# Cortical tracking of voice pitch in the presence of multiple speakers depends on selective attention

**DOI:** 10.1101/2021.12.03.471122

**Authors:** Christian Brodbeck, Jonathan Z. Simon

## Abstract

Voice pitch carries linguistic as well as non-linguistic information. Previous studies have described cortical tracking of voice pitch in clean speech, with responses reflecting both pitch strength and pitch value. However, pitch is also a powerful cue for auditory stream segregation, especially when competing streams have pitch differing in fundamental frequency, as is the case when multiple speakers talk simultaneously. We therefore investigated how cortical speech pitch tracking is affected in the presence of a second, task-irrelevant speaker. We analyzed human magnetoencephalography (MEG) responses to continuous narrative speech, presented either as a single talker in a quiet background, or as a two-talker mixture of a male and a female speaker. In clean speech, voice pitch was associated with a right-dominant response, peaking at a latency of around 100 ms, consistent with previous EEG and ECoG results. The response tracked both the presence of pitch as well as the relative value of the speaker’s fundamental frequency. In the two-talker mixture, pitch of the attended speaker was tracked bilaterally, regardless of whether or not there was simultaneously present pitch in the speech of the irrelevant speaker. Pitch tracking for the irrelevant speaker was reduced: only the right hemisphere still significantly tracked pitch of the unattended speaker, and only during intervals in which no pitch was present in the attended talker’s speech. Taken together, these results suggest that pitch-based segregation of multiple speakers, at least as measured by macroscopic cortical tracking, is not entirely automatic but strongly dependent on selective attention.

## Introduction

Pitch is a function of temporal periodicity and spectral order in acoustic waveforms (de Cheveigné, 2005). The cochlea transforms temporal periodicity into a spatial code by mapping different frequencies in the signal to different spatial locations along the basilar membrane. Subcortical responses retain the periodicity in ranges critical for speech, and thus represent pitch temporally as well as spatially (Skoe and Kraus, 2010; Maddox and Lee, 2018). However, phase locking at faster frequencies gradually declines in the ascending auditory system (Joris et al., 2004). Extracranial recordings of cortical responses have observed population-level phaselocking to periodicity in speech only up to approximately 110 Hz (Coffey et al., 2016; Kulasingham et al., 2020). This is insufficient for encoding most voice pitch, as voices often exhibit a fundamental frequency above 100 Hz. Instead, the auditory cortex is abundant with frequency-selective receptive fields (Saenz and Langers, 2014), and pitch features are encoded through a combination of place and rate code (Fishman et al., 2013).

Cortical pitch tracking has primarily been analyzed with higher level representations of pitch. In speech, pitch is present intermittently, forming the basis of voiced segments and interrupted for unvoiced segments. Two aspects of pitch can thus be described separately that are relevant for cortical tracking: *pitch strength*, i.e., to what extent pitch is present at each moment in the speech signal, and *pitch value*, i.e., the height of the perceived pitch, generally corresponding to the fundamental frequency. Both these features are tracked by scalp electroencephalography (EEG) responses to continuous narrative speech (Teoh et al., 2019). Intracranial recordings suggest that representations of relative pitch, corresponding to speaker-independent intonation contours, are more prominent than representations of absolute pitch (Tang et al., 2017); and that the pitch of speech is associated with a prominent neural response at around 100 ms latency (Li et al., 2021).

Here we investigate how pitch tracking is affected when listening to multiple simultaneous speakers. When the sound from two speakers is mixed, the sound waveforms combine additively. For simplicity we will consider the case of a single audio channel mixed signal presented diotically, i.e., the two source waveforms are mixed into a single mixed waveform presented to both ears. The problem of stream segregation then is segregating the spectro-temporal elements of the heard sound into those associated with either of the sources (Bregman, 1990). Pitch can be a strong cue for stream segregation (Bregman, 1990; Micheyl and Oxenham, 2010). For example, pitch tracking can aid segregation by grouping together the different harmonics of a shared fundamental (Popham et al., 2018). The spatial code in A1 provides sufficient information to distinguish two concurrent vowels that differ in f0 by four semitones, consistent with human perceptual judgements (Fishman et al., 2014, 2016). Non-primary areas might thus reconstruct pitch from this representation (Bendor and Wang, 2005), for example using harmonic templates (Fishman et al., 2014). This would potentially allow the auditory cortex to segregate the pitch of two speakers, especially if those two streams differ substantially in pitch (e.g., a male and a female speaker). A pitch sensitive region in the anterior portion of the superior temporal plane (Norman-Haignere et al., 2013) could be the potential locus for such pitch-based segregation.

If pitch extraction is automatic for each of several multiple sources in a mixture, it could then be used as bottom-up cue in stream segregation. This would be consistent with suggestions that the subcortical representation of voice pitch (Maddox and Lee, 2018; Van Canneyt et al., 2021a, 2021b) is affected by attention (Forte et al., 2017; Etard et al., 2019; Saiz-Alía et al., 2019). Cortical responses might then be expected to simultaneously track the pitch in the attended and the ignored speakers. Pitch tracking might still be affected by overlapping pitch to the extent that the overlap imposes additional demands for segregation. On the other hand, pitch tracking might reflect a secondary representation constructed during attentive speech processing, for example for linguistic prosody. In this case, pitch tracking might depend on selective attention, possibly without demonstrating pitch tracking for the ignored speaker at all.

To investigate this, we analyzed a previously studied dataset of MEG responses to audiobooks in two conditions: speech from a single speaker in a quiet background, and speech from two speakers, one male and one female, reading different audiobooks mixed together and presented diotically, with the task of listening to one speaker and ignoring the other (Brodbeck et al., 2020). In the original study we analyzed responses as a function of spectrogram representations, and found that listeners segregate acoustic features even of the ignored speaker from the acoustic mixture. Here we ask to what degree listeners additionally track pitch in the attended and the ignored speaker. In this analysis of pitch tracking, all predictors used in the original analysis are also controlled for (Brodbeck et al., 2020). For clean speech, we model pitch through two separate time-dependent predictors, pitch strength and pitch value (**Figure 1**-A). For the two-speaker mixture, we additionally distinguish (1) between pitch in the attended and the ignored talker, and (2) between pitch when it is *overt*, i.e., when only one of the two speakers exhibits pitch at a time, and when it is *masked*, i.e., when both speech signals contain pitch simultaneously (**Figure 1**-B). Masked pitch deserves special attention because the two sources of pitch may interfere with each other, such that naïve pitch detection algorithms would fail without considering the effect of that interference through some kind of segregation mechanism (Micheyl and Oxenham, 2010).

**Figure 1.**
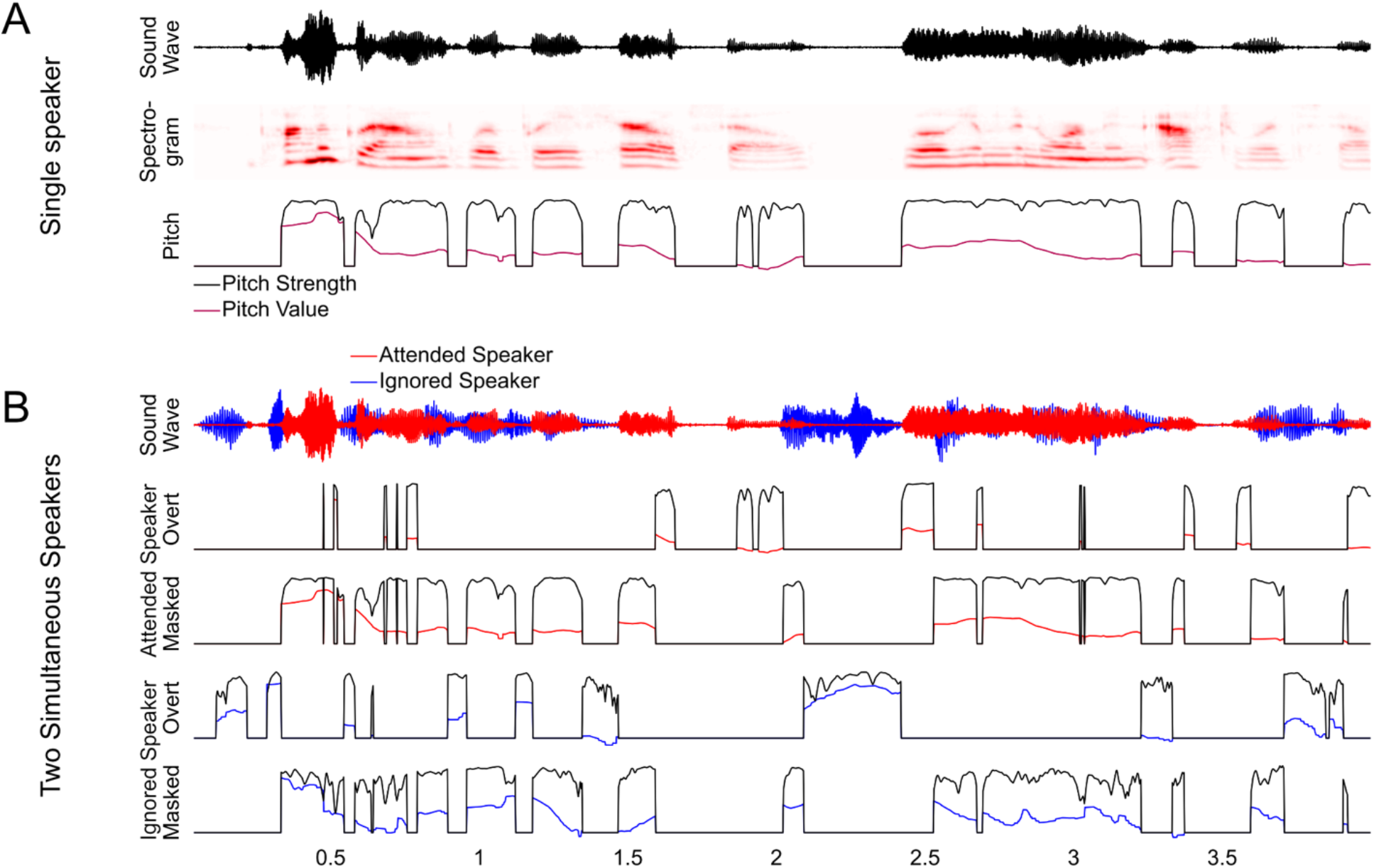
Predictors for analyzing pitch tracking. **(A)** For a single speaker, pitch tracking was estimated using two predictors: *Pitch Strength*, reflecting the degree to which a distinctive pitch is present in the sound signal, and *Pitch Value*, reflecting the fundamental frequency of the pitch, relative to baseline. For moments when Pitch Strength is 0, Pitch Value is set to the default baseline value. **(B)** For two-speaker stimuli, pitch strength and value were estimated separately for each speaker, and then split into two separate predictors reflecting overt pitch (i.e., pitch is present only in a single speaker) and masked pitch (i.e., pitch is present in both speakers). Note that, as a consequence of this definition, the two masked pitch predictors are always on simultaneously, whereas the overt pitch predictors are mutually exclusive.

## Method

We reanalyzed MEG responses from 26 native speakers of English (8 female, 18 male; age mean 45.2 years, range 22-61), listening to multiple 1 minute duration audiobook segments, in quiet and in a two-talker mixture (8 and 16 minutes, respectively). Mixtures always consisted of a male and a female speaker, with clearly separable by pitch. Most of the analysis followed essentially the same procedures as the original study (Brodbeck et al., 2020) but with additional predictors to isolate representations of pitch.

MEG recordings were pre-processed in mne-python (Gramfort et al., 2014) with temporal signal-space separation (Taulu and Simola, 2006), a 1-40 Hz bandpass filter, independent component analysis (Bell and Sejnowski, 1995), and an additional 20 Hz lowpass filter. Responses were then projected to current dipoles oriented perpendicularly to the white matter surface (4-fold icosahedral subdivision) using distributed minimum *ℓ*2 norm source current estimates.

### Predictors

Pitch was extracted from each stimulus using Praat (Boersma and Weenink, 2017). Pitch strength was taken directly from the Praat estimate. The pitch value was log transformed, and sections without pitch (pitch strength of zero) were set to the 5^th^ percentile value of sections with pitch (see **Figure 1**-A). This baseline correction was performed for each 1-minute segment separately, to derive relative pitch regardless of the specific speaker’s fundamental frequency. The pitch value was oriented relative to the lower end of the scale to account for the observation in ECoG that native English speakers show selective responses to higher relative pitch (Li et al., 2021). To control for spectro-temporal acoustic processing we included the acoustic spectrogram and onset spectrogram predictors from the original study (8 bands each).

We also considered a pitch onset predictor (Krumbholz et al., 2003), based on the half-wave rectified derivative of the pitch strength. We reasoned that this predictor might be able to isolate responses related to the initial detection of pitch. However, the predictor did not improve predictive power beyond pitch strength and value (*t_max_*=2.28, *p*=.385, when restricted to the STG), and we consequently dropped it from further analysis. A reason for this might be that pitch onset in speech almost always coincides with a sound onset in the spectrogram, which our analysis always controlled for.

For the two-speaker condition, we first generated pitch predictors for each of the two source segments in the mixture. Masking was operationalized as a binary distinction: The ignored speaker was considered masked where the pitch strength of the attended speaker exceeded 0.5 and vice versa. Based on this, both speaker’s pitch predictors were split into two different sets, one reflecting overt pitch, the other masked pitch (**Figure 1**-B). To control for spectro-temporal processing we included all predictors from the two talker condition of the original study, including overt and masked onsets (Brodbeck et al., 2020, second equation on p. 17).

### Model tests

Multivariate temporal response function (mTRF) models were estimated separately for each subject and source dipole with Eelbrain (Brodbeck et al., 2021). As in the original study, models with latency range 0-500 ms were estimated and tested on held-out data using 4-fold crossvalidation. Predictive power was quantified as the proportion of the variability in the source localized MEG responses explained by the model. Each predictor was evaluated by comparing the predictive power of the complete model (all predictors) with a model that was estimated while excluding the to-be-tested predictor.

We defined anatomical areas for mass-univariate tests (based on “aparc” labels; Desikan et al., 2006): For pitch representations of clean speech we initially tested in the whole cortex with the exception of the occipital lobe, insula and cingulate cortex (i.e., regions in which we did not expect a substantive auditory response, excluded in order to expedite these numerically intensive computations). Based on these results, we performed tests for the two-speaker condition in more restricted areas in the superior temporal gyrus (STG; transverse and superior temporal gyrus labels) and the inferior frontal gyrus (IFG; pars opercularis, pars triangularis and pars orbitalis labels). Anatomical maps of predictive power were smoothed (SD = 5 mm) and compared with mass-univariate related measures *t*-tests, correcting for multiple comparison with threshold-free cluster enhancement (Smith and Nichols, 2009) and a permutation distribution based on 10000 random permutations of condition labels. Tests of whether a given predictor improved predictive power were one-tailed, all other comparisons were two-tailed. Even though we sometimes report results separately for the left and right hemisphere, multiple comparison correction was always based on a permutation distribution estimated from the combination of both hemispheres.

To express model predictive power as a meaningful quantity, the predictive power of different predictors is expressed as % of the explanatory power of the most complete model (separately for the single speaker and the two-speaker conditions).

There is no standard measure of effect size for mass-univariate tests. As a compromise, we report *t*_max_ for mass-univariate tests, i.e., the largest t-value in the significant area (or the whole tested area for non-significant results). However, to provide a more traditional measure of effect size, we also defined a function region of interest (ROI). This ROI was defined based on the intersection of significant activation in the single speaker condition (to either of the two pitch predictors) and the STG anatomical area. We used this ROI to extract the average explained variability attributable to pitch strength and value combined (**Figure 3**-B) or each predictor individually (**Table 1**).

**Table 1.** In the two-talker mixture, pitch tracking is dominantly due to pitch value, not pitch strength. Each row shows the unique predictive contribution of one predictor, in the two-speaker condition, in an STG ROI based on significant activity in the one speaker condition. Shown are a one-sample *t*-test of the difference in prediction accuracy when excluding a given predictor, and Cohen’s *d*.

| Speaker | Masking | Pitch | Left Hemisphere |  |  |  | Right Hemisphere |  |  |  |
| --- | --- | --- | --- | --- | --- | --- | --- | --- | --- | --- |
|  |  |  | *t* | *p* |  | *d* | *t* | *p* |  | *d* |
| Attended | Overt | Strength | 2.09 | .023 | * | 0.41 | 0.36 | .361 |  | 0.07 |
|  |  | Value | 4.21 | < .001 | *** | 0.83 | 3.18 | .002 | ** | 0.62 |
|  | Masked | Strength | 0.85 | .202 |  | 0.17 | -1.44 | .919 |  | -0.28 |
|  |  | Value | 3.85 | < .001 | *** | 0.75 | 3.45 | .001 | ** | 0.68 |
| Ignored | Overt | Strength | 0.71 | .242 |  | 0.14 | -0.25 | .600 |  | -0.05 |
|  |  | Value | 1.76 | .045 | * | 0.35 | 2.90 | .004 | ** | 0.57 |

### ANOVA for difference in localization

Localization *differences* in MEG should be interpreted with caution (Lütkenhöner, 2003; Bourguignon et al., 2018). However, the question whether two localizations are based on *the same* underlying source configuration can be tested in a straightforward manner, based on the linearity of the forward and inverse models. If two maps reflect the same underlying source configuration, the should produce the same relative measurements at the sensor level (McCarthy and Wood, 1985) and, consequently, in the source localized responses. Based on this, we test the null hypothesis that two predictors are represented in the same neural sources by first normalizing the two respective maps and subtracting one form the other. If the two underlying maps have the same shape, the sources should now only contain random noise. We thus used a one-way repeated measures ANOVA with factor source dipole to test whether there is a systematic pattern left after the subtraction (in other words, whether there is a systematic difference between the patterns of localization of the two predictors).

### Temporal response functions

To analyze temporal response functions (TRFs), mTRF models were re-estimated using a latency range from −100 – 500 ms, and without held-out data (but still using early stopping based on cross-validation). For TRF estimation, predictors as well as MEG responses were normalized, and TRFs were analyzed at this normalized scale.

## Results

### Pitch strength and value are tracked in single talker speech

In single talker clean speech, pitch strength and pitch value were both represented neurally (**Figure 2**). Both predictors separately improved prediction of brain responses, over and above auditory spectrogram representations (strength: *t_max_*=5.58, *p*<.001; value: *t_max_*=6.00, *p*<.001). Overall, source localization is consistent with the majority of sources in the vicinity of the auditory cortex in Heschl’s gyrus and the superior temporal gyrus (see **Figure 2**-A). Pitch strength was significantly right-lateralized (Figure 2-B; *t_max_*=4.90, *p*<.001), with no significant tracking in the left hemisphere (*t_max_*=3.24, *p*=.670).

**Figure 2.**
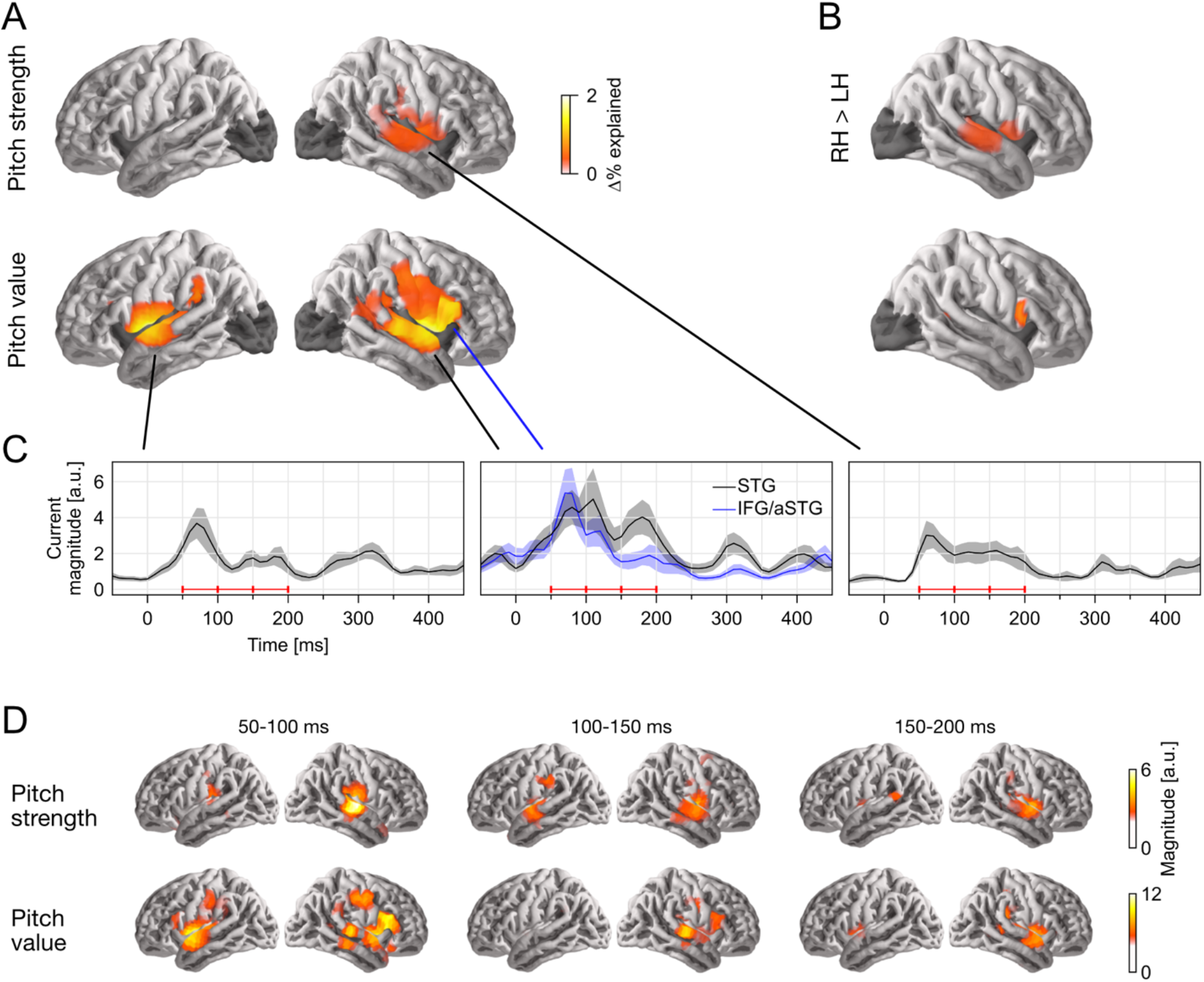
Separable tracking of pitch strength and pitch value of a single talker. **(A)** Pitch strength and pitch value both improved model predictions independently, when controlling for acoustic envelope and onset spectrograms (*p* ≤ 0.05, corrected; darkened areas excluded from analysis). The color scale reflects the explained variability in MEG responses, expressed as % of the complete model. **(B)** Both pitch predictors showed some right-lateralization. Plots show right–left hemisphere predictive power difference, same scale as (A). **(C)** Temporal response functions (TRFs) showed dominant responses at latencies between 50-200 ms. TRF magnitude is shown for regions of significant model prediction. The three horizontal red bars indicate time windows used in (D). **(D)** Anatomical distribution of TRFs in 50 ms time windows. LH: left hemisphere; RH: right hemisphere; STG: superior temporal gyrus; IFG: inferior frontal gyrus; aSTG: anterior STG.

The localization of pitch value was more complex: when tested in the whole brain, it was significantly right-lateralized (*t_max_*=4.90, *p*<.001). However, the region of significant difference coincided with the anatomical label of the pars opercularis of the inferior frontal gyrus (IFG). When repeated in the STG only, lateralization was not significant (*t_max_* = 2.28, *p* = .385). To confirm that pitch strength and value are tracked by non-identical sources we applied a predictor × source dipole ANOVA in the right hemisphere (see Methods). This indicated that the distribution of sources tracking the two predictors was indeed different (*F*(117, 2925) = 1.85, *p* < .001). Together, these results suggest that pitch value tracking engages additional, more anterior sources compared to pitch strength tracking. The source localization raises the possibility that pitch value specifically engages the right IFG, although due to the proximity to the anterior temporal lobe it is impossible to exclude the possibility of an anterior temporal source with dispersion into IFG due to imperfect source localization (cf. Bourguignon et al., 2018).

### Response to pitch peaks around 100 ms latency

The Temporal Response Functions (TRFs), i.e., the estimated impulse responses to elementary pitch features, and are shown in in **Figure 2**-C and D. **Figure 2**-C shows the response magnitude, summed across source dipoles, as a function of time. Responses are shown in functional ROIs, based on combining the region of significant model predictions (union across the two predictors) with anatomical STG and IFG labels. Most of the response power is concentrated in the first 50-200 ms, with a clear response peak to pitch value around 100 ms. Comparison of responses to pitch value in the STG and IFG ROIs suggests that the relative involvement of the anterior peak is stronger at the shorter latencies. The anatomical distribution of the response magnitude is consistent with this, showing a stronger response at the anterior source in the early time window (**Figure 2**-D).

### In two simultaneous talkers, pitch-tracking depends on selective attention

To test how pitch is tracked when listening to one of two concurrent talkers, we generated 4 versions of each predictor: first, we generated separate versions for pitch of the attended speaker and of the ignored speaker; second, for each of those, we separated each time point into overt or masked pitch, based on whether pitch was simultaneously present in the other talker or not (see **Figure 1**-B). First, we tested for pitch tracking in the STG by combining pitch strength and value in each of the four categories (**Figure 3**-A). Results indicated significant pitch tracking for overt pitch, regardless of whether pitch originated from the attended (*t_max_*=4.38, *p*<.001) or the ignored speaker (*t_max_*=3.74, *p*=.008). In contrast to this, masked pitch was tracked only in the attended speaker (*t_max_*=3.57, *p*=.003), whereas we did not find evidence for tracking of masked pitch in the unattended speaker (*t_max_*=2.42, *p*=.432). A direct comparison in the STG confirmed that masked pitch in the attended speaker was tracked significantly stronger than in the unattended speaker (*t_max_*=3.88, *p*=.005). Although tracking of overt pitch in the ignored speaker was not significant in the left hemisphere, the lateralization was not significant either (*t_max_*=2.47, *p*=.054). Nevertheless, comparing overt pitch tracking in the attended and the ignored speaker suggested a small effect of selective attention (*t_max_*=3.26, *p*=.031), with stronger tracking of the attended speaker in the left hemisphere only, not in the right (*t_max_*=2.48, *p*=.415). Only attended masked pitch was also significant in the IFG/aSTG area (*t_max_*=3.20, *p*=.025), and in this area attended masked pitch was significantly stronger than ignored masked pitch (*t_max_*=3.54, *p*=.017) as well as attended overt pitch (*t_max_*=3.40, *p*=.035).

**Figure 3.**
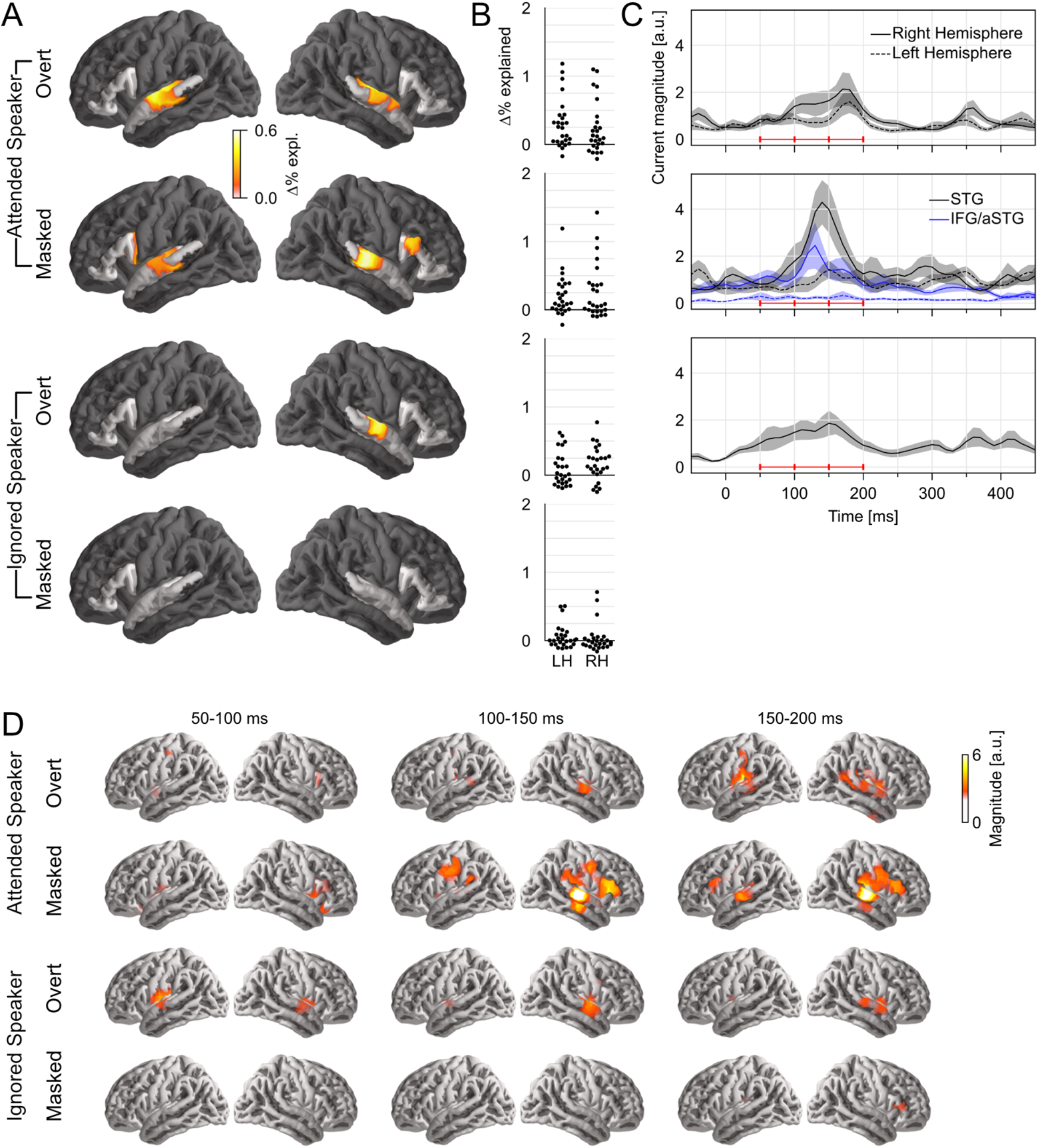
Pitch tracking in two simultaneous speakers depends on selective attention. **(A)** Significance tests of pitch tracking for overt and masked pitch in the attended and ignored speaker. STG and IFG were separately tested (darkened area excluded from tests). **(B)** Individual subject data (% variability explained) in a region of interest, defined as the intersection of the region of significant activity in the single speaker condition and the STG anatomical label. **(C)** Temporal response function (TRF) magnitude with dominant response at 100-200 ms latency. The three horizontal red bars indicate time windows used in (D). **(D)** TRF activity localized mainly to the auditory cortex, with involvement of a more anterior region for masked pitch in the attended speaker. LH: left hemisphere; RH: right hemisphere; STG: superior temporal gyrus; IFG: inferior frontal gyrus; aSTG: anterior STG.

Next, we asked whether pitch strength and pitch value were independently tracked for concurrent speakers in each of the significant categories. Surprisingly, in mass-univariate tests in the STG none of the pitch strength predictors were significant, while all three pitch value predictors were. To derive proper measures of effect size we also performed univariate tests in a ROI based on significant STG activation in the one speaker condition (**Table 1**). While these results are largely consistent with the mass-univariate tests, they do suggest marginally significant tracking of overt pitch strength in the left hemisphere. The univariate tests are likely less conservative than the mass-univariate tests, by not correcting for multiple comparisons in the two hemispheres. Nevertheless, the result suggests that some pitch strength tracking might exist, although with a much weaker effect size than pitch value tracking.

Based on the above results, we only analyzed TRFs to pitch value. The TRFs to overt pitch in the attended and the ignored speaker were qualitatively similar (**Figure 3**-B, C). In contrast, masked pitch in the attended speaker was associated with a large peak in the right STG around 140 ms.

## Discussion

Our analysis of responses to clean speech confirms a previous report of separate cortical pitch strength and pitch value tracking seen with EEG (Teoh et al., 2019). In addition, source localization suggested a differentiation between the two representations, with a right-lateralized STG representation of pitch strength and a bilateral STG representation of pitch value, with potential additional involvement of a more anterior region. While source localization shows a more anterior localization in IFG, we cannot exclude the possibility of this being an artifact of imperfect source localization (Bourguignon et al., 2018), as a source in anterior STG would be more consistent with fMRI reports of a pitch representation in the anterior STG (Norman-Haignere et al., 2013).

In the presence of two simultaneous speakers, pitch tracking depends on selective attention, but not exclusively. Overt pitch was similarly represented, regardless of whether that pitch was in the attended or the ignored speaker. This suggests that overt pitch extraction occurs without a need for selective attention, and might form part of an auditory background representation. On the other hand, when pitch was present in both speakers simultaneously, selective attention had a strong effect: pitch in the attended speaker was tracked very robustly, with recruitment of additional, more anterior neural sources, possibly reflecting additional resources recruited for speaker segregation. At the same time, we found no evidence for a representation of pitch in the ignored speaker.

These results have implications for the functional relevance of cortical pitch tracking. Cortical voice pitch tracking might reflect purely acoustic processing, i.e., the extraction of pitch and the pitch trajectory (e.g. Andermann et al., 2021). However, pitch also carries linguistic meaning. Even in non-tonal languages (like English), the pitch contour is a prosodic cue that carries information on the information structure and phrase structure of an utterance. Pitch tracking could thus also reflect processing of such higher-level properties. Our observation of overt pitch tracking in the ignored speaker suggests that pitch tracking at least does not exclusively reflect higher-level processing, given that linguistic processing of ignored speech in such situations is very limited (Brodbeck et al., 2018; Broderick et al., 2018).

### Stream segregation of a monoraul mixture is cortical and depends on selective attention

A long-standing question on cocktail party speech processing is whether segregation of multiple speakers occurs pre-attentively, with selective attention merely selecting one of multiple input streams, or post-attentively, with selective attention actively contributing to the segregation. Recent evidence support the latter view, at least when the speech signals are mixed together monophonically, i.e. without spatial separation cues (Puvvada and Simon, 2017; O’Sullivan et al., 2019; Brodbeck et al., 2020). Our new results are consistent with this. On the one hand, significant tracking of overt pitch in the ignored speaker suggests that pitch tracking itself does not require selective attention, as long as the pitch is easily extracted through the periodicity of the signal. However, in masked pitch we found a strong effect of selective attention, with no evidence of tracking of ignored pitch at all. Consistent with the previous reports, the present results do not provide evidence for pre-attentive pitch-based segregation, but do suggest enhanced pitch processing in a selectively attended speaker.

### Lateralization

Cortical pitch processing has sometimes been specifically associated with the right hemisphere. For example, pitch judgements engage the right prefrontal cortex (Zatorre et al., 1992), and the right auditory cortex might play a causal role in pitch discrimination learning (Matsushita et al., 2021). Our results provide additional evidence for a tendency towards right-lateralization of at least some aspects of pitch tracking in speech. However, while we found some evidence for stronger pitch representations of clean speech in the right hemisphere, we did not find any evidence of lateralization during the two-speaker condition, suggesting that pitch processing might become more bilateral in the more demanding condition.

### Conclusion

The central finding of this study is that cortical pitch tracking is modulated by selective attention. While listeners represent overt pitch similarly in an attended or an ignored speaker, they do not seem to track pitch of an ignored speaker that is masked by pitch in the attended speaker. In contrast, tracking of masked pitch is robust for an attended speaker, suggesting that this pitch is selectively extracted and processed.

## Acknowledgements

This work was supported by the National Institutes of Health (R01-DC014085 and R01-DC019394 for JZS) and the National Science Foundation (SMA-1734892 for JZS and 1754284 for CB [J. Magnuson, PI]).

